# The Plasmepsin-Piperaquine Paradox Persists

**DOI:** 10.1101/2024.11.28.625831

**Authors:** Breanna Walsh, Robert L Summers, Dyann F Wirth, Selina Bopp

## Abstract

Malaria is still a major health issue in many parts of the world, particularly in tropical and subtropical regions of Africa, Asia, and Latin America. Despite significant efforts to control and eliminate the disease, malaria remains a leading cause of illness and death, mainly due to the occurrence of drug-resistant parasites to the frontline antimalarials such as dihydroartemisinin-piperaquine (DHA-PPQ). Artemisinin resistance has been linked to kelch13 mutations, while decreased PPQ sensitivity has been associated with higher *plasmepsin II* and *III* gene copies and mutations in the chloroquine resistance transporter.

In this study, we demonstrate the effective use of CRISPR/Cas9 technology to generate single knockouts (KO) of *plasmepsin II* and *plasmepsin III*, as well as a double KOs of both genes, in two isogenic lines of Cambodian parasites with varying numbers of plasmepsin gene copies. The deletion of *plasmepsin II* and/or *III* increased the parasites’ sensitivity to PPQ, evaluated by the area under the curve. We explored several hypotheses to understand how an increased *plasmepsin* gene copy number might influence parasite survival under high PPQ pressure. Our findings indicate that protease inhibitors have a minimal impact on parasite susceptibility to PPQ. Additionally, parasites with higher *plasmepsin* gene copy numbers did not exhibit significantly increased hemoglobin digestion, nor did they produce different amounts of free heme following PPQ treatment compared to wildtype parasites. Interestingly, hemoglobin digestion was slowed in parasites with *plasmepsin II* deletions. By treating parasites with digestive vacuole (DV) function modulators, we found that changes in DV pH potentially affect their response to PPQ. Our research highlights the crucial role of increased *plasmepsin II and III* gene copy numbers in modulating response to PPQ and begins to uncover the molecular and physiological mechanisms underlying PPQ resistance in Cambodian parasites.

**Author Summary:** Global malaria control has plateaued, with drug-resistant *Plasmodium falciparum* posing a significant challenge. Artemisinin-based combination therapies (ACTs) are becoming less effective, especially in South-East Asia, where resistance to dihydroartemisinin-piperaquine (DHA-PPQ) is leading to treatment failures, notably in Cambodia. Genome-wide studies link artemisinin resistance to kelch13 mutations, while decreased PPQ sensitivity is tied to higher *plasmepsin II* and *III* gene copies and mutations in chloroquine resistance transporter.

We previously showed a connection between increased plasmepsin gene copies and reduced PPQ sensitivity. In this study we try to understand the biological role of the plasmepsins in PPQ sensitivity. Therefore, we knocked out *plasmepsin II* and *III* genes in Cambodian strains using CRISPR/Cas9, and found increased PPQ sensitivity, confirming these genes’ roles in resistance. Plasmepsins are proteases that participate in the hemoglobin degradation cascade in the digestive vacuole of the parasites. Protease inhibitor experiments and hemoglobin digestion studies indicate that digestive vacuole pH fluctuations affect PPQ response, highlighting the need for further research into PPQ resistance mechanisms.

## Introduction

Despite remarkable strides towards global malaria control in the past two decades, progress in reducing the total number of malaria cases has plateaued (1). Drug resistance in *Plasmodium falciparum* parasites, particularly to first- and second-line artemisinin (ART) combination therapies (ACTs), remains a major challenge to the elimination of malaria. In regions of South-East Asia (SEA), resistance to both potent ART-derived compounds and long-lasting partner drugs is emerging and rapidly spreading (2). Widespread parasite resistance to dihydroartemisinin-piperaquine (DHA-PPQ) has resulted in clinical treatment failures throughout western Cambodia, where DHA-PPQ has been used since 2008 (3–7). PPQ resistance is marked by parasite recrudescence 42 days post-treatment (3). The spread of PPQ-resistant parasites in SEA has significantly shortened the usable lifespan of DHA-PPQ, resulting in Cambodia’s 2016 reversal to artesunate-mefloquine at a first-line antimalarial (8).

Genome-wide association studies (GWAS) of parasites collected from patient samples in SEA through the Tracking Resistance to Artemisinin Collaboration (TRAC) identified mutations in the propeller region of *kelch13* as highly linked with ART treatment failure (9, 10). GWAS of Cambodian *P. falciparum* isolates have revealed that decreased PPQ sensitivity is associated with increased copy numbers of the genes coding for the aspartic proteases *plasmepsin II* and *III* in combination with a single copy of the multidrug resistance-1 (*pfmdr1*) gene (11, 12). In vitro studies in the laboratory strain 3D7 identified a slight sensitization when *plasmepsin II* and *III* were deleted (13) but no susceptibility change was observed when *plasmepsin II* and *III* were overexpressed (14). Furthermore, SNPs in the chloroquine resistance transporter (*pfcrt*) have also been associated with PPQ resistance (5, 11, 12, 15). Mutations in *pfcrt* have been shown to confer PPQ resistance when introduced into the Dd2 laboratory strain (16–19). To date, parasites with mutations in *kelch13*, increased *plasmepsin* copy numbers, and *pfcrt* mutations are now almost fixed in SEA (20–23), the treatment failure rate is up to 70% in Western Cambodia (2) and surveillance of these markers is now underway in Africa and South America.

Despite the identification of promising molecular markers, the characterization of PPQ resistance in parasite field isolates has proved challenging. When subjected to a standard drug susceptibility assay, resistant parasites yield bimodal dose-response curves, with increased parasite survival at high PPQ concentration. These bimodal curves cannot be described using traditional non-linear regression analysis and yield non-interpretable EC_50_ values when a curve fit is forced (4, 7). As a result, many practitioners rely on estimates of the EC_90_ instead. A PPQ survival assay (PSA) was developed, wherein parasite survival after 48h of a single PPQ treatment (200nM) is compared to that of parasites cultured in vehicle control (5). However, PSA analysis requires manual slide counting, making it is both labor and time intensive. To better describe the bimodal response, we increased the concentration range of PPQ and utilized the Area Under the Curve (AUC) of the high-dose peak to quantify the PPQ response (24). Culture adapted TRAC isolates showed a correlation between increased *plasmepsin II* and *plasmepsin III* copy numbers and AUC. A panel of clonal isogenic lines, all with single copies of *pfmdr1* and identical *pfcrt* loci, showed decreased sensitivity to PPQ with increasing copy numbers of *plasmepsin II* and *plasmepsin III*, implicating these copy number variations (CNVs) as potential drivers of PPQ resistance (24). Analysis of *plasmepsin II* and *plasmepsin III* knock outs (KOs) in the relevant genetic background of Cambodian isolates is key to further understanding the role of genetic background in modulating Plasmepsin II and Plasmepsin III action under PPQ pressure.

Plasmepsin II and III, along with Plasmepsin I and IV, are aspartic proteases that participate in the hemoglobin degradation cascade in the digestive vacuole (DV) (25–29). Targeted genetic disruptions of the *plasmepsin* genes-either individually or in combination-yield viable parasites lacking dramatic changes in morphology or growth, suggesting functional redundancy between these aspartic proteases and other proteases in the DV (29, 30). The digestion of hemoglobin crucially provides a source of amino acids for parasites; however, the breakdown of hemoglobin releases toxic free heme, which aggregates as inert, crystalline hemozoin within the DV (31–33). As an aminoquinoline, PPQ is thought to impede the degradation of hemoglobin within the parasite DV, leading to a build-up of toxic free heme and parasite death (17, 33). However, the biological mode of PPQ action is not well understood.

Here, we show successful CRISPR/Cas9-mediated *plasmepsin II* single KO, *plasmepsin III* single KO, and *plasmepsin II* and *plasmepsin III* double KO in two isogenic lines of Cambodian parasites with variable *plasmepsin* copy numbers. Disruption of *plasmepsin II* and/or *plasmepsin III* resulted in increased parasite sensitivity to PPQ, as measured by the AUC. We tested several hypotheses as to how increased *plasmepsin* copy number could influence survival under high PPQ pressure. We show that protease inhibitors have a minimal effect on parasite susceptibility to PPQ. In addition, hemoglobin digestion was not significantly increased in parasites with higher plasmepsin copy numbers nor did these parasites produce different amounts of free heme upon PPQ treatment. However, hemoglobin digestion was slowed in parasites with *plasmepsin II* deletions. To explore how the physiological conditions of the DV contribute to PPQ susceptibility, we treated parasites with modulators of DV function, suggesting that fluctuations in DV pH affect parasite response to PPQ. Thus, in Cambodian parasites, we describe the critical role of *plasmepsin II* and *plasmepsin III* copy number variations in modulating parasite response to PPQ and begin to probe the molecular and physiological underpinnings of PPQ resistance.

## Results

### Generation of plasmepsin KO in TRAC isolates

We have previously shown that clones from the TRAC isolate KH001_053 contained variations in *plasmepsin II* and *III* copy numbers that correlate with their resistance phenotype, as measured by the AUC in PPQ growth assays. These clones are genetically identical, including the *Pf*crt (Dd2 like) and the *Pf*mdr1 loci (single copy); the only difference is the plasmepsin II/III CNV (24). We confirmed the tandem arrangement of the duplicated *plasmepsin II* and *III* locus suggested by Amato et al. (12, 34), where the break points of the duplication are in the 3’ regions of *plasmepsin I* and *plasmepsin III*, resulting in a chimeric *plasmepsin III/I* between two *plasmepsin II* copies, followed by an intact *plasmepsin III* copy (Fig 1A). To determine if either *plasmepsin II* or *plasmepsin III* is responsible for the increased AUC in PPQ growth assays, we generated *plasmepsin II* and *plasmepsin III* single knockouts (KOs) and a *plasmepsin II/III* double KO in the relevant genetic background of a TRAC isolate. A PPQ-susceptible clone of KH001_053 with a single copy of *plasmepsin II* and *III* (KH001_053_G10_) and a PPQ-resistant clone with two copies of both *plasmepsin II* and *plasmepsin III* (KH001_053_G8_) served as parental lines (24). We used CRISPR/CAS9 technology to introduce single strand breaks in the *plasmepsin* genes and provided the parasites with a template to disrupt either *plasmepsin II* (G10_PMII_KO_ and G8_PMII_KO_) and or *plasmepsin III* (G10_PMIII_KO_ and G8_PMIII_KO_) alone or both genes simultaneously (G10_PMII/III_KO_ and G8_PMII/III_KO_) with an *hdhfr* selectable marker as described previously (29). Gene editing in the duplicated KH001_053_G8_ locus resulted in the same outcome as the KH001_053_G10_ single locus due to CAS9 cutting both copies of the duplicated *plasmepsin* genes and fusion of the two remaining pieces with the *hdhfr* marker (See Fig 1 for the cloning strategy).

**Figure 1.**
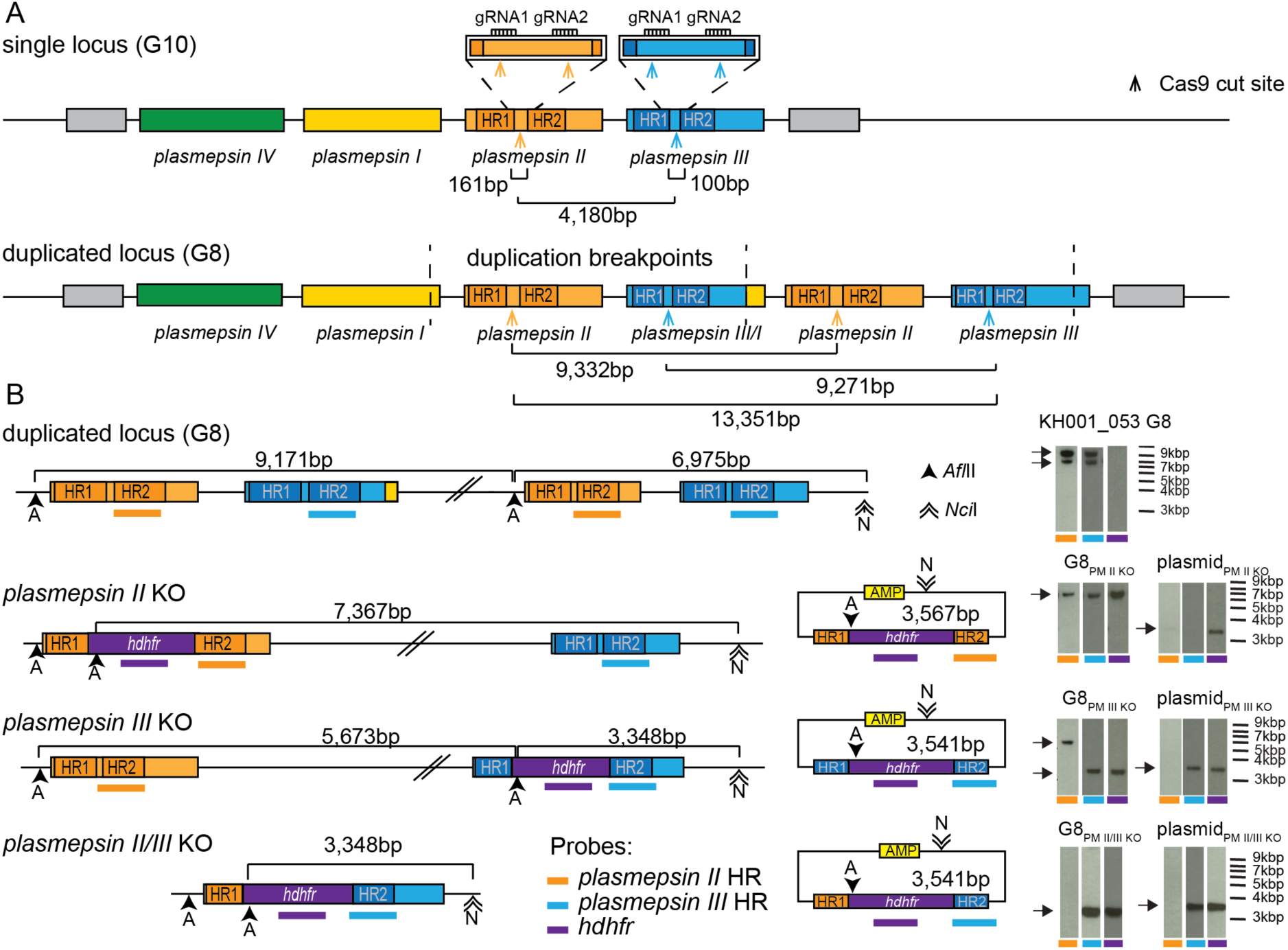
Correct disruption of the duplicated plasmepsin locus by KO constructs. A) Schema of the single and duplicated plasmepsin loci predicted by Amato et al. (12). B) Schema of duplicated locus, edited loci and homology plasmids used for integration into the locus. Restriction enzyme sites and expected band sizes for Southern blots are indicated in the schema, gDNA was digested [*Afl*II (A) and *Nci*I (N)], run on a gel, transferred to a membrane, and hybridized with three different probes indicated by colored bars (orange: *plasmepsin II*, blue: *plasmepsin III* and purple: *hdhfr* cassette). Arrows indicate expected band sizes. Southern blots for additional clones and the single locus parasites (KH001_053_G10_) are shown in S1-3 Fig.

Transgenic parasites were cloned, and subsequently, the integration status was verified by PCR, quantitative PCR (qPCR) (S1 Table), and Southern blots for at least two clones per transfection (Fig 1 and S1-3 Fig).

### Loss of plasmepsin duplication in field isolates abolishes PPQ bimodal response

We tested the KH001_053_G10_ and KH001_053_G8_ parental lines and the engineered *plasmepsin* KO parasites in PPQ growth assays, compared the initial hill slope, and measured the AUC (Fig 2 A-E and S2 Table). It should be noted that the KO parasites generated from KH001_053_G8_ reduced the *plasmepsin* copy numbers from two copies, and the KH001_053_G10_ KO parasites from one copy, resulting in genetically identical KOs. Consequently, the engineered parasites from KH001_053_G10_ and KH001_053_G8_ were also indistinguishable in their phenotypic response (Fig 2E, one-way ANOVA with Tukey post-test p<0.05). We therefor used KH001_053_G8_ KO lines for further phenotypic analysis.

**Figure 2.**
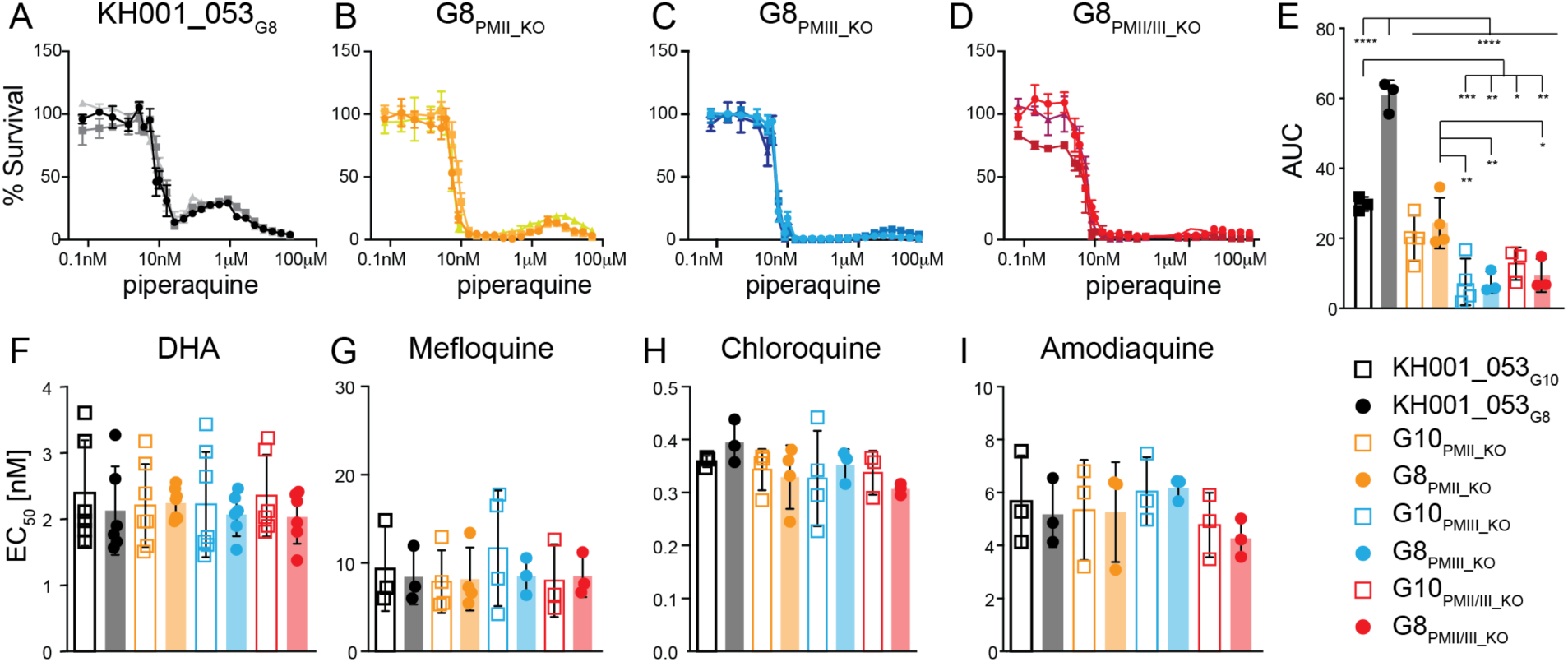
Disruption of plasmepsin loci leads to loss of AUC. Parasites were exposed to increasing levels of PPQ for 84h and survival was measured by increased DNA content. Shown are an example of three biologically independent experiments run in triplicates for the KH001_053_G8_ parental locus (A) and parasites with disruption of either *plasmepsin II* (G8_PM II_KO_, B), *plasmepsin III* (G8_PM III_KO_, C), or a double knockout of *plasmepsin II/III* (G8_PM II/III_KO_, D). The area under the curve (AUC) between the local minima was calculated and the average and SD is shown in for KH00_053_G10_ and KH001_053_G8_ parents as well as KOs (E). Statistics show one-way ANOVA with Tukey post-test: *, p < 0.05; **, p < 0.01; ***, p < 0.001; ****, p < 0.0001. Standard EC_50_ values for the PPQ partner drug DHA (F), and three PPQ analogues (Mefloquine (G), Chloroquine (H), and Amodiaquine (I)) are shown as average and SD for at least three biological replicates run in triplicates. The two parental lines and their relative plasmepsin KO lines are shown. No statistically significant differences were detected between the lines.

G8_PMII_KO_, G8_PMIII_KO_, and G8_PMII/III_KO_ had indistinguishable initial hill slopes but had a strong significant reduction of the AUC when compared to the KH001_053_G8_ parent with two copies (Fig 2A-D, one-way ANOVA with Tukey post-test p<0.0001). The same was true for G10_PMII_KO_, G10_PMIII_KO_, and G10_PMII/III_KO_ compared to KH001_053_G8_. However, only G8_PMIII_KO_, G8_PMII/III_KO_, G10_PMIII_KO_, and G10_PMII/III_KO_ showed significantly reduced AUC compared to the KH001_053_G10_ single locus (Fig 2E and S2 Table, one-way ANOVA with Tukey post-test p<0.05). Taken together, these data confirm the role of duplicated *plasmepsin* genes in survival of parasites at high concentration of PPQ and *Plasmepsin III* appears to be the dominant driver of this phenotype.

### Loss of plasmepsins does not affect susceptibility to other antimalarials

To understand if *plasmepsin* copy number variations modulate susceptibility to other antimalarials, we also subjected the engineered *plasmepsin* KO lines to a panel of related quinolines and DHA (i.e., the partner drug of PPQ in ACTs) in standard *in vitro* drug susceptibility assays. There were no significant differences in EC_50_ between the parental lines and the clonal *plasmepsin* KO lines for chloroquine (CQ), mefloquine (MEF), amodiaquine (AQ) or DHA (Fig 2F-I, S2 Table; one-way ANOVA with Tukey posttest, p>0.05). As expected for parasites with a Dd2-like *Pfcrt*, the parental lines and *plasmepsin* KO lines were all resistant to CQ compared to the CQ-sensitive laboratory strain 3D7 (EC_50_: 12nM, S2 Table). These data suggest that the role of *plasmepsin* copy numbers in increased survival is specific to PPQ and does not extend to other closely related quinolines.

### Aspartic and cysteine protease inhibitors do not impact the AUC

Given that deletion of *plasmepsin II* and *plasmepsin III* abolished the PPQ-resistance phenotype, we wondered if direct inhibition of the protease function of the Plasmepsins might reduce the AUC. We demonstrated that protease inhibitors E64 (a broad-band cysteine protease inhibitor shown to inhibit falcipain-2 in the DV (35) and host egress (36)) and Pep A (shown to inhibit plasmepsin function in cell lysates and bind to *plasmepsin II* (37)) had no differential activity in parasites regardless of plasmepsin copy number (S2 Table).

We next investigated if there was an additive effect of the protease inhibitors in combination with different PPQ concentrations in parasites with increased *plasmepsin* copy numbers. We chose two different drug concentrations where parasite growth was affected but not severely inhibited (69-89% growth, S4 Fig) and kept them constant for each protease inhibitor (7.5 and 5μM for pepA methyl ester and 2.6 and 3.9μM for E64) in the presence of increasing concentrations of PPQ. We tested the parental lines with the single and the duplicated locus as well as an additional TRAC isolate with an even higher AUC (KH004_057, AUC=88±21 (24)). The tested protease inhibitor concentrations had no significant effect on the AUC (p > 0.05, one-way ANOVA followed by Tukey posttest), suggesting that disruption of hemoglobin catabolism by protease inhibitors does not modulate PPQ susceptibility under the conditions tested here (S5 Fig, S3 Table).

### PPQ induced heme accumulation is not impacted by *plasmepsin II or III* copy numbers

To further test the possible mechanism of PPQ resistance, we tested the effect of PPQ on hemozoin biocrystallization using a pyridine-labeled heme fractionation assay (38, 39). This assay uses a series of cellular fractionation steps to extract the different heme species in the parasites (hemoglobin, free heme, and hemozoin) and subsequently measures the Fe(III)heme-pyridine absorbance. It has been demonstrated previously that parasites exposed to increasing CQ or PPQ concentrations increase the level of free heme and hemoglobin while reducing the levels of hemozoin (17).

To understand if the additional copies of *plasmepsin II* and *III* influence the generation of free heme or the degradation of hemoglobin under PPQ pressure, we exposed highly synchronized ring stage parasites (0-6h) from the KH001_053_G8_ and KH001_053_G10_ parental lines to a range of PPQ concentrations for 32h and collected the different heme species. We determined the percentage of each heme species present in the total amount of iron extracted. On average, both parental lines showed similar percentages of all heme species in the absence of drug (13% free heme, 4% hemoglobin and 83% hemozoin, Fig 3A-C, S4 Table). As shown previously (17), parasites exposed to 200nM or 2μM PPQ showed a significant increase in free heme and hemoglobin compared to the untreated control, as well as a reduction in hemozoin formation (unpaired Student’s t test p < 0.05). These findings are consistent with an inhibition of hemoglobin degradation and hemozoin formation by PPQ treatment. There was no significant difference between the parental lines containing one (KH001_053_G10_) or two (KH001_053_G10_) copies of *plasmepsin II/III* in any of the heme species. This suggests that while both parasite lines experience similar levels of the toxic free heme, parasites with increased plasmepsin copy numbers seem to be better adapted to survive these extremely high concentrations of PPQ.

**Figure 3.**
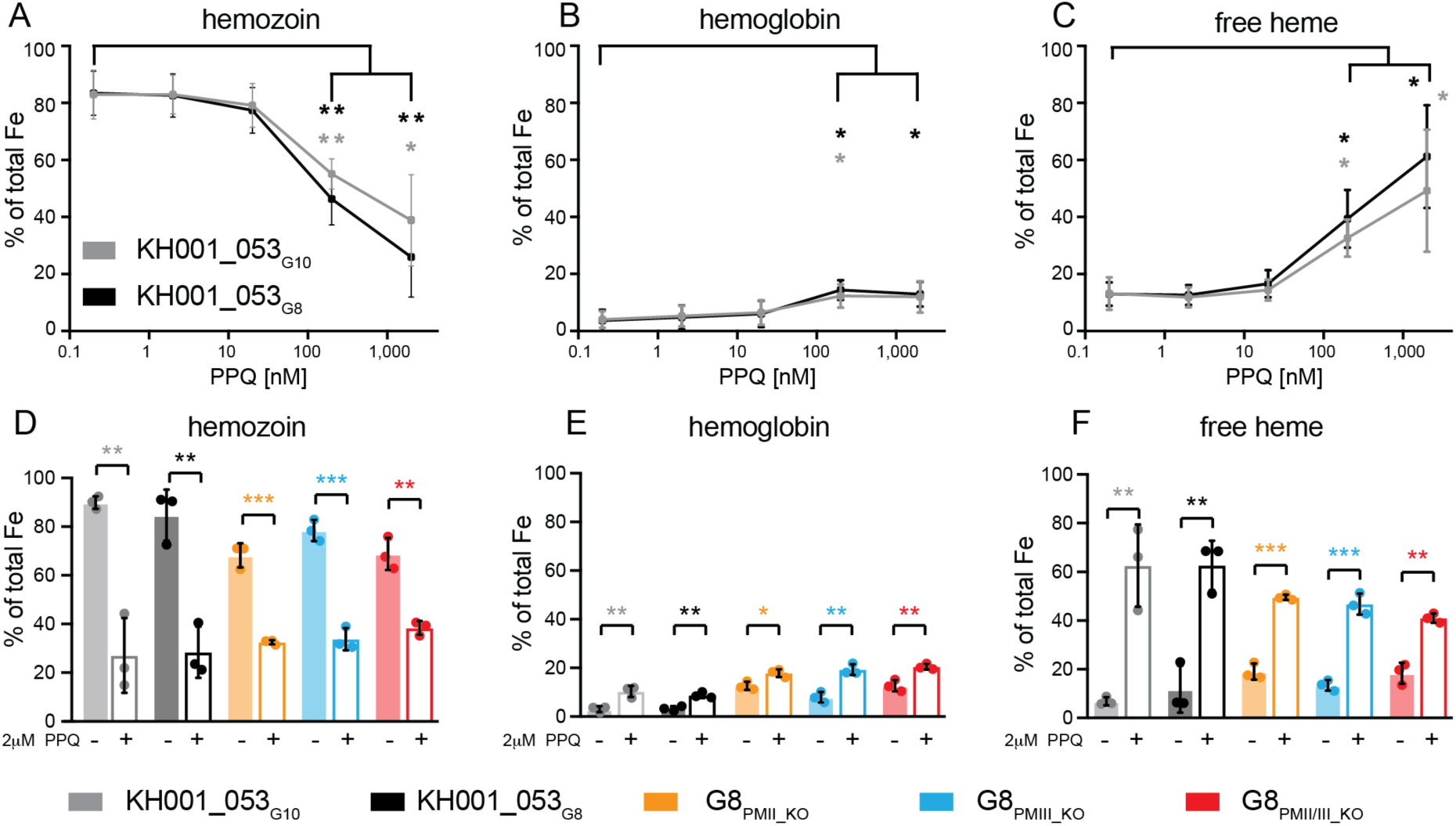
Heme fractionation of PPQ treated and untreated parasites with variable plasmepsin copy numbers. Different heme species were extracted from parasites by subsequent cellular fractionation steps. A-C: Tightly synchronized KH001_053_G10_ (grey) and KH001_053_G10_ (black) ring stage parasites were exposed to various PPQ concentrations for 32h. Average and SD of percentage of hemozoin Fe (A), hemoglobin (B), and free heme Fe (C) are shown for three independent experiments run in quadruplicates. Statistical comparisons of the drug-treated lines to their untreated controls were performed using two-tailed unpaired student’s t tests *, p < 0.05; **, p < 0.01. D-F: KH001_053_G10_ and KH001_053_G8_ parental lines as well as G8_pmII_KO_, G8_pmIII_KO_, and G8_pmII/III_KO_ were incubated with 2μM PPQ from 24 to 36h post synchronization and harvested at 36h. Statistical comparisons of treated to untreated parasites at 36h timepoint from three independent biological replicates were performed using two-tailed unpaired student’s t tests *, p < 0.05; **, p < 0.01, p < 0.01, *** p > 0.001.

We next exposed parental and KH001_053_G8_ KO parasites (G8_PMII_KO_, G8_PMIII_KO_, and G8_PMII/III_KO_) to a single dose of 2μM PPQ for 12h between the 24 and 36h post invasion when hemoglobin digestion is highest. As seen before, even a 12h exposure of parasites to 2μM PPQ lead to a significant decrease in hemozoin, and an increase in free heme and hemoglobin compared to 36- 42h untreated control in all lines tested (Fig 3D-F and S5 Table, unpaired student’s t test, p< 0.05). However, free heme accumulation was higher in the parental lines (>60%) than in the KO lines (<50%), suggesting that less plasmepsin activity leads to less free heme. This demonstrates that PPQ is inhibiting hemozoin formation regardless of plasmepsin abundance or presence.

### Hemoglobin digestion is slowed in plasmepsin KO parasites

To further understand the role of the plasmepsins in hemoglobin degeneration, we investigated if parasites with increased copy numbers or deletions of *plasmepsin II* and *III* differ in their composition or accumulation of heme species during their life cycle. We tightly synchronized the two parental lines KH001_053_G10_ and KH001_053_G8_ as well the KO clones G8_PMII_KO_, G8_PMIII_KO_, and G8_PMII/III_KO_ and collected timepoints 24-, 36-, and 42h post-synchronization (Fig 4). As expected, the hemozoin percentage increased over time for all lines, while hemoglobin and free heme concentrations decreased (Figure 4A-C and S5 Table). Throughout the lifecycle, there was significantly less hemozoin contribution in G8_PMII_KO_, and G8_PMII/III_KO_ lines compared to the KH001_053_G10_ single copy line (Fig 4A, unpaired student’s t test p < 0.05). In accordance, hemoglobin and free heme levels were significantly higher in the G8_PMII_KO_ and G8_PMII/III_KO_ lines compared to KH001_053_G10_, suggesting that hemoglobin digestion is slowed in the absence of *plasmepsin II.* G8_PMIII_KO_ parasites showed an intermediate phenotype between the single copy parent KH001_053_G10_ line and *the* G8_PMII_KO_ and G8_PMII/III_KO_ lines (Fig 4B and C). A similar increase in undigested hemoglobin has been observed in PPQ resistant parasite lines carrying the *Pf*CRT G353V or F145I variants (18) as well as in *plasmepsin II* KOs detected by Western blot (30). There was no difference between the two parental lines, suggesting that hemoglobin digestion efficiency is not increased in parasites with additional plasmepsin copy numbers. There was no statistically significant difference in heme species between the time point at 36 and 42h (by unpaired student’s t test) indicating that most of the hemoglobin digestion is completed by 36- 42h. The reduced hemoglobin digestion efficiency in KO parasites could explain the observation that less free heme is released under PPQ treatment (Fig 3F).

**Figure 4.**
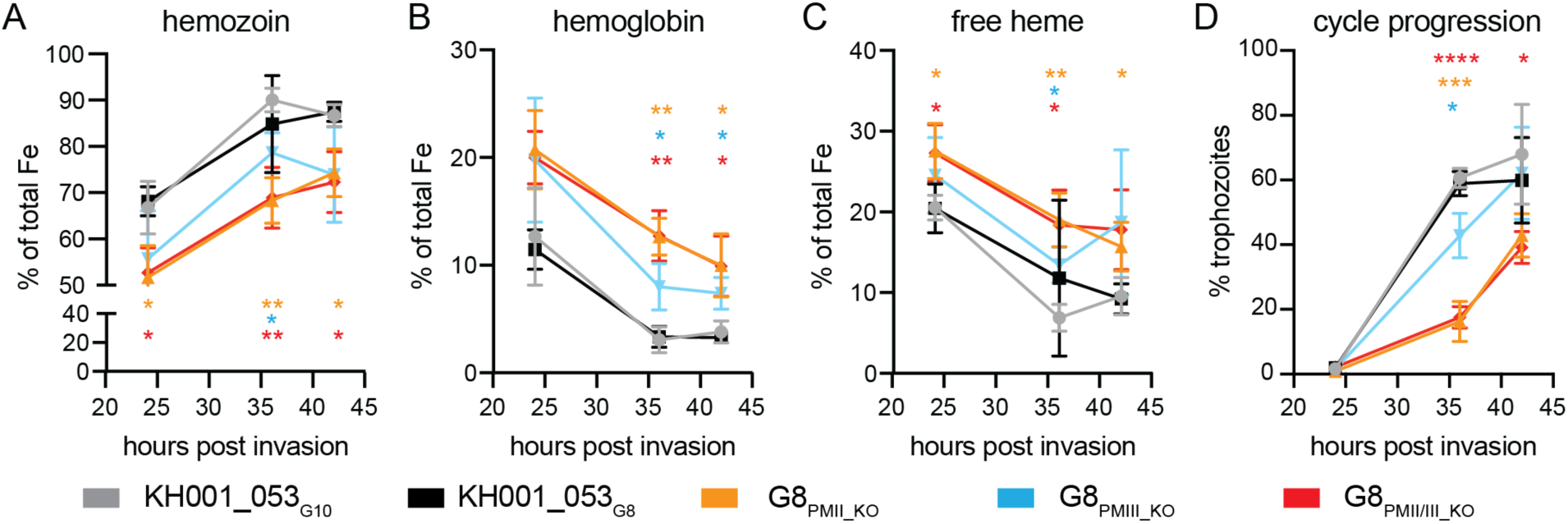
Plasmepsin KO parasites have slowed hemoglobin metabolism. Tightly synchronized parasites were harvested at different time-points throughout the life cycle and average and SD of percentage of hemozoin Fe (A), hemoglobin (B), and free heme Fe (C) are shown for three independent experiments for parasites with a single locus (KH001_053_G10_ grey), duplicated locus (and KH001_053_G8_ black), G8_pmII_KO_ (orange), G8_pmIII_KO_ (blue), or G8_pmII/III_KO_ (red). Statistical comparisons at each time point were performed between the single copy KH001_053_G10_ parasite and all other lines using two-tailed unpaired student’s t tests *, p < 0.05; **, p < 0.01. D) Cell cycle progression was measured by the number of nuclei present per cell using flow cytometry and SYBRGreen staining of DNA. Parasites were defined as trophozoites when they had more than three nuclei. Shown is the percentage of trophozoites at each time point. Statistical comparisons were performed between the single copy KH001_053_G10_ parasite and all other lines at the same time point using two-tailed unpaired student’s t tests *, p < 0.05; ***, p < 0.001, ****, p < 0.0001.

We next asked if the delay in hemozoin formation seen in G8_PMII_KO_ and G8_PMII/III_KO_ is correlated to a delay in overall life cycle progression. To measure cell cycle progression, we determined the percentage of trophozoites in each sample by FACS flow prior to the iron species extraction (Fig 4D). In agreement with the iron species data, trophozoite formation was delayed in the *plasmepsin* KO parasites compared both KH001_053_G10_ and KH001_053_G8_ parental lines, with the G8_PMII_KO_, and G8_PMII/III_KO_ lines being more delayed than the G8_PMIII_KO_ line. It remains unclear if the delay in hemoglobin digestion is the result or the cause of slowed progression through the life cycle.

### PPQ activity is not dependent on external pH

Drug uptake and availability to the parasite can be affected by the drugs protonation status and its ability to cross membranes. Changes in the pH of the extracellular environment have been shown to affect CQ potency (40, 41). CQ is membrane permeable at neutral pH, but loses its permeability once protonated in the low pH of the DV (42) and concentrates within the DV via ‘weak-base trapping’ and subsequent binding to ferriprotoporphyrin IX (43). Decreasing the pH of the extracellular environment reduces the overall pH gradient between the DV and extracellular environment and leads to reduced CQ accumulation in the DV and increased survival of the parasites. Indeed, in media with a pH adjusted to 6.74, EC_50_ values for CQ dramatically increased when compared to their EC_50_ values at neutral pH of 7.5 (S6A-F Fig and S6 Table). Moreover, Dd2, KH001_53_G10_, and KH001_53_G10_ were not completely killed at the highest CQ concentration tested (2μM). Increasing the pH of the media to 8.24 had the opposite effect, significantly reducing the EC_50_ of CQ when compared to media at a pH of 7.5 (p>0.01, paired Student t-test, S6 Fig and S6 Table). This is consistent with a larger pH gradient between the extracellular medium (at pH 8.24) and the DV leading to increased CQ accumulation within the DV.

PPQ, like CQ, is a weak base, and we hypothesized that its protonation could be affected by the pH of its surrounding medium. In *in vitro* PPQ susceptibility assays, parasites showed only marginal higher sensitivity to high PPQ concentrations at increased extracellular pH, and the differences in AUC between the different pH conditions were not statistically significant (Fig 5A-D, S6 Table). This suggests that the activity of PPQ is less affected by external pH than the CQ activity.

**Figure 5.**
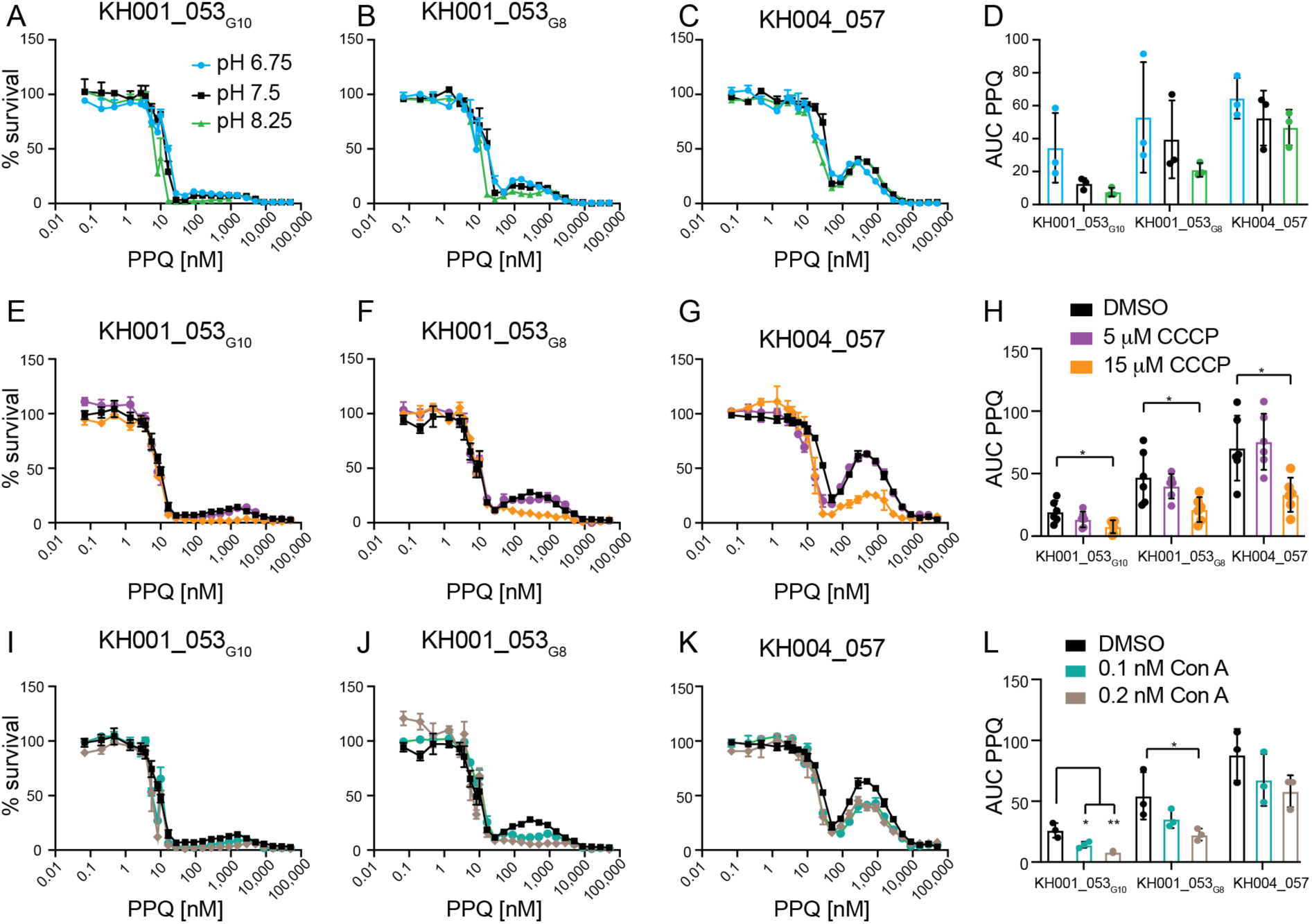
Effect of pH on PPQ AUC. A-D: Parasites were exposed to increasing levels of PPQ for 84h in acidic (pH6.74), normal (pH7.5) or basic (pH8.24) media. Shown are an example of three biologically independent experiments run in triplicates for KH001_053_G10_ (A), KH001_053_G8_ (B) or KH004_057 (C), as well as the average and SD of the AUC for all three biological replicates (D). There were no statistically significant differences detected. E-L: Parasites were exposed to increasing levels of PPQ in the presence of DMSO or E-H) conA at either 0.1nM or 0.2nM or I-L) CCCP at either 5 μM or 15μM concentrations. Shown are one example of three biologically independent experiments run in triplicates for KH001_53_G10_ (E, I), KH001_53_G8_ (F, J), and KH004_057(G, K). The area under the curve (AUC) between the local minima was calculated and the average and SD are shown for conA (H) and CCCP (L). Statistical comparisons of the drug-treated lines and DMSO-treated controls were performed using one-way ANOVA with Dunnett’s post-test: *, P < 0.05 and **, P < 0.01.

### Alkalization of the DV affects PPQ AUC

The hemoglobin digestion activity of DV lysates (27, 44), as well as recombinant plasmepsins (25, 26), is maximal at low pH, consistent with conditions within the acidic DV (DV pH of about 4.5-5.5 (45–47)). We therefore reasoned that perturbations to the DV pH might also impact plasmepsin activity and thereby alter the PPQ-resistance phenotype of parasites with increased *plasmepsin II and III* copy numbers. We used concanamycin A (conA, an inhibitor that blocks the V-type ATPase from pumping H^+^ ions into the DV lumen, thereby leading to an alkalization of the DV (48)) and carbonyl cyanide m-chlorophenyl hydrazone (CCCP, a proton ionophore dissipating H^+^ gradients across membranes in general) to interrogate the role of the DV pH in PPQ resistance.

We first determined the EC_50_ for conA and CCCP. While we did not determine any differences in susceptibility to conA, G8_PMII_KO_ and G8_PMII/III_KO_ parasites were less susceptible to CCCP than KH001_053_G10_ (p < 0.05, one-way ANOVA followed by Dunnett posttest, S3 Table). Previous work demonstrated that vacuolar pH impacted CQ efficacy (41, 49), and we similarly found a significant increase in susceptibility to CQ when we tested CQ in the presence of CCCP at 5μM or 15μM concentration against parasites lines with various plasmepsin copy numbers (p < 0.01, one-way ANOVA followed by Tukey posttest, S6G-L Fig and S3 Table). Interestingly, the decreased survival was specific for the CQ resistant *Pf*CRT isoform and had non-significant effect on the sensitive 3D7 parasite line.

Similarly, when treated with 15μM CCCP, parasites tested in an PPQ susceptibility assay exhibited reduced AUC values compared to parasites not treated with CCCP, indicating that intracellular pH also affects the PPQ resistant phenotype (p < 0.05 one-way ANOVA followed by Dunnett posttest, Fig 5E-H). When we tested conA in combination with PPQ, we also found a significant reduction in the AUC for KH001_053_G10_ and KH001_053_G8_ but not for KH004_057 (Fig 5I-L). We next wanted to test whether PPQ itself could affect the alkalization of the DV.

### PPQ does not influence digestive vacuole pH

The weak-base trapping effect of CQ can cause alkalinization of acidic compartments (50). The activities of plasmepsin enzymes are pH-sensitive and are most active at low pH. Hence, we hypothesized that if PPQ also acts as a weak base accumulator, sufficiently high concentrations of PPQ in the DV might disrupt the DV pH, thereby reducing Plasmepsin activity, and the production of free heme which leads to parasite death. Amplification of *plasmepsin II* and *III* could therefore rescue this reduced activity due to increased protein expression, leading to the survival of PPQ-resistant parasites under high PPQ concentrations. To test this hypothesis, we used ratiometric fluorescence measurement of Dd2 trophozoite parasites with fluorescein-dextran loaded DVs to monitor DV pH in the presence and absence of PPQ. Fluorescence traces for pH calibration buffers indicated resting DV pH values of 5.5 ± 0.2 (mean ± SD, n = 4; Fig 6), consistent with previous studies (46, 51). The V-type ATPase inhibitor Con A (100 nM), proton ionophore CCCP (10 µM), and weak-base NH_4_Cl (10 mM) all caused rapid alkalinization of parasite DVs, resulting in significantly elevated DV pH compared to the vehicle control following 45 minutes of exposure (6.5-6.7; p<0.05, One-way ANOVA with Dunnett correction for multiple comparisons, Fig 6 and S7 Table). CQ at 10 µM also caused significant alkalinization, resulting in an average DV pH of 6.4 ± 0.2 (p < 0.05, One-way ANOVA, Dunnett post-test). By contrast, concentrations of up to 50 µM of PPQ had no impact on DV pH (Fig 6), suggesting that PPQ does not exert an effect via pH modulation over this time-course. This provides further evidence that PPQ and CQ have distinct effects on DV physiology.

**Figure 6.**
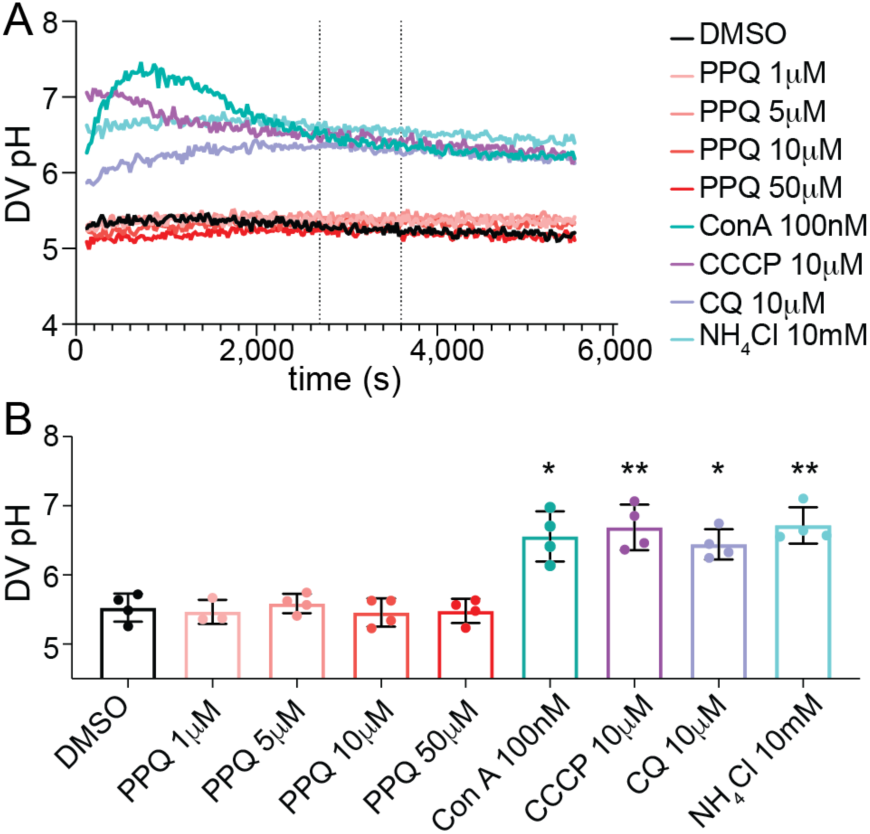
Digestive vacuole pH measurement of fluorescein-dextran loaded Dd2 parasites. A) DV pH traces of Dd2 parasites exposed to PPQ and control treatments Concanamycin A (ConA, 100 nM), CCCP (10 µM), CQ (10 µM) and NH_4_Cl (10 mM). Shown are the averaged results of internal technical duplicates from a representative experiment. DV pH was quantified for each treatment as an average of the measurements taken between 45 and 60 mins of compound exposure (dashed vertical lines), B) average DV pH measurements following 45-60 minutes of exposure to PPQ at 1, 5, 10 or 50 µM and to known DV pH modulators Concanamycin A, CQ, CCCP and NH_4_Cl. The data are the average and SD of 3 to 4 independent experiments (performed with blood from different donors). The asterisks denote a significant difference from the DMSO solvent control: ** *P* < 0.01, * *P* < 0.05 (one-way ANOVA). Raw data are provided in S7 Table.

## Discussion

We explored the contribution of plasmepsins copy numbers specifically in the second peak of the bimodal resistance phenotype to PPQ measured by the AUC in the absence of *pf*crt mutations. We generated *plasmepsin II, III,* and double KO parasites in the relevant genetic background of Cambodian isolates. Resulting plasmepsin KO parasite lines had lost their secondary survival peak measured by the AUC clearly demonstrating that plasmepsins are the main driver of the AUC phenotype.

We investigated several potential biochemical and metabolic mechanisms underlying the AUC phenotype. However, we were unable to define a specific mechanism explaining the genetic results. We demonstrated decreased hemoglobin metabolism in plasmepsin KO parasites, consistent with the role of plasmepsins in hemoglobin digestion (25–29). However, multiple copies of *plasmepsin II* and *III* did not increase hemoglobin digestion and therefore the AUC cannot be explained by altered hemoglobin digestion. Analysis of pH changes in the DV demonstrated that PPQ had no effect on DV pH in contrast to proton trapping observed with CQ by us and others (50). Additionally, perturbance of the parasite’s proton gradient had only minimal effect on parasite’s PPQ susceptibility, again in contrast to a hyper sensitization to CQ in the presence of CCCP. Therefore, PPQ AUC resistance is unlikely to be mediated by vacuolar pH.

Previous work, both in natural isolates (2, 5, 15, 20) and engineered parasites (16, 17, 23, 52), has demonstrated that additional mutations in the Dd2-like *Pf*CRT (C101F, T93S, H97Y, F145I, I218F, M343L, and G353V) mediate PPQ resistance in the absence of plasmepsin copy number changes. Functional studies showed that *Pf*CRT is capable of transporting PPQ, and PPQ resistance conferring mutations in *Pf*CRT increase PPQ transport (53, 54). In the experiments presented here, the role of *Pf*CRT was controlled as all lines used did have the Dd2-like haplotype (M74I, N75E, K76T, A220S, Q271E, N326S, I356T, and R371I) and therefore the changes in the AUC phenotype cannot be explained by *Pf*CRT.

However, there is the possibility that the increase in plasmepsin copy numbers is a compensatory variation to the combination of DHA and PPQ treatment in ACTs. The parasites used in this experiment all had a K13 C580Y mutation. K13 mutations have been shown to cause lower hemoglobin uptake and digestion in early ring stages (55, 56). An increase in *plasmepsin* copy number might provide a compensatory function of increased hemoglobin digestion in early ring stages. Unfortunately, we were unable to reliably measure hemoglobin digestion in early ring stages and therefore could not directly test this hypothesis. Indeed, a positive association between K13 C580Y and increased plasmepsin copy numbers was observed in a genetic cross between a wildtype Malawian parasite and a Cambodian line (57). Further evidence of the benefit of increased plasmepsin copy numbers comes from genetic data in field isolates in SEA where *plasmepsin* amplification occurred on a large diversity of haplotypes independent of DHA-PPQ selection pressure (22). Increased copy numbers of *plasmepsins* have been described in 2007 for clinical isolates with K13 mutations (20, 58), while mutations in *Pfcrt* were only reported after 2010 (59) but got to almost fixation in 2016 on an increased plasmepsin copy number background in SEA (16). In Africa, increased *plasmepsin II* copy numbers have been detected in parasites collected between 2014-2015, especially in Burkina Faso and Uganda (> 30%)(60). It is also possible that the peptide composition or accumulation from hemoglobin digestion is different in parasites with higher plasmepsin copy numbers and further investigation is needed to explore this possibility (18).

Previous work by others has demonstrated that plasmepsins are not essential for parasite survival under *in vitro* conditions with *plasmepsin III* disruption being more severe (29, 31, 61, 62). Similar to our results here, these studies also detected only minimal changes in susceptibility of the KO lines to protease inhibitors E64 or pepA (29, 30).

Two genetic crosses with a sensitive parasite and a PPQ resistant TRAC parasite harboring increased plasmepsin copy number and a mutation in *Pf*crt have been generated and analyzed for their PPQ phenotypes (57, 63). While the major contributor to high PSA survival was mapped to *Pfc*rt, higher copy numbers of *plasmepsin II/III* increased the resistance even further. When AUC was used as a phenotype for QTL, the main peak was still *Pfc*rt but a second strong peak was found around plasmepsins suggesting a link between *Pf*CRT and plasmepsins (63).

Overall, in the natural setting, plasmepsin copy number variation has a role in conferring PPQ resistance. Here we demonstrate that the AUC phenotype, namely increased survival under high PPQ pressure, is mediated by plasmepsin copy number. While we demonstrate the importance of plasmepsins in mediating increased AUC in resistant parasites we still don’t understand the biological role of plasmepsins in PPQ resistance.

## Materials and Methods

### Parasite Culture

The parental parasite lines (KH001_053 and KH004_057) were collected from Pursat and Pailin, Cambodia in 2011 through the Tracking Resistance to Artemisinin Collaboration (TRAC), and culture adapted and subcloned as previously described resulting in KH001_053_G10_ and KH001_053_G8_ (24). All parasites were grown in fresh human erythrocytes (O+) at 4-5% hematocrit in Roswell Park Memorial Institute (RPMI) 1640 media supplemented with 26.6 mM NaHCO_3_, 27.7 mM Hepes, 0.41 mM hypoxanthine, 10% O+ human serum, and 25 ug/mL gentamicin. Human serum (heat inactivated and pooled) and human erythrocytes were supplied by Interstate Blood Bank, Inc., Memphis, TN. For *in vitro* drug susceptibility assays, parasites were grown in media containing 0.5% Albumax (Invitrogen), in place of human serum. Cultures were incubated at 37°C with rotation (55 RPM) under 1% O_2_/5% CO_2_/balance N_2_ gas.

### Design and Construction of CRISPR/Cas9 Plasmid Vectors for Transfection

Transfections were performed using a two-plasmid strategy. The first plasmid contained homology regions to the gene of interest (GOI) flanking a *hdhfr* positive selectable marker, while the second plasmid contained the coding sequence for Cas9 and a guide RNA (gRNA) specific to the GOI.

To generate the basic template vector pGEM*hdhfr*+, the *hdhfr* positive selectable marker cassette was PCR amplified from the vector pL-6_eGFP(64) with primers *hdhfr* 5’ F and *hdhfr* 3’ R and cloned into pGEM-3Z (Promega, Madison, WI) digested with *Hinc*II. Two flanking regions corresponding to the GOI were then cloned into this basic vector to generate a template for homologous recombination and disruption of the GOI. The homology regions were amplified from 3D7 gDNA with the following primer sets (adapted from Liu et al., (29)) - *plasmepsin II* 5’ homology region: PMII 5’ HR Fwd and PMII 5’ HR Rev; *plasmepsin II* 3’ homology region: PMII 3’ HR Fwd and PMII 3’ HR Rev; *plasmepsin III* 5’ homology region: PMIII 5’ HR Fwd and PMIII 5’ HR Rev; and *plasmepsin III* 3’ homology region: PMIII 3’ HR Fwd and PMIII 3’ HR Rev. The 3’ homology regions were digested with *Pst*I and *Sph*I and ligated into the appropriately digested pGEM*hdfr+* basic template vector. The coordinating 5’ homology regions were digested with *Afl*II and *Xma*I and ligated into the pGEM*hdfr+* vector already containing the 3’ homology region. The *plasmepsin II/III* double KOs construct contains a *plasmepsin II* 5’ HR and a *plasmepsin III* 3’ HR flanking the *hdhfr* cassette. The homology region plasmids were transfected into parasites in either their circular or *Bgl*I-linearized form. GOI-specific gRNA sequences were generated using Benchling (https://benchling.com) and ligated into the *Bbs*I digested pDC2-Cas9-U6-hDHFR vector(65). All primers used are described in S8 Table.

### Parasite Transfection and Selection

Parental KH001_053_G8_ and KH001_053_G10_ parasites were synchronized using 5% D-sorbitol (Sigma-Aldrich, St. Louis, MO) and early ring stage parasites were mixed with 50 μg of each transfection plasmid in cytomix (120 mM KCl, 0.15 mM CaCl2, 2 mM EGTA, 5 mM MgCl2, 10 mM K2HPO4/KH2PO4, pH 7.6, 25 mM Hepes, pH 7.6) and electroporated in 2mm cuvettes at 0.31kV and 950uF (66). Electroporated cells were then placed in O^+^ media with fresh erythrocytes at 5% hematocrit. After 24-48h of drug-free recovery, cultures were continuously treated with 5 nM WR99210, which positively selects for parasites containing *hdhfr*. Recovered parasites were genotyped as bulk cultures and subcloned by limiting dilution. Subsequently, clonal lines were genotyped by PCR, qPCR, and Southern blotting. All primers used are described in S8 Table.

### Polymerase Chain Reaction (PCR) for Genotyping Transfectants

Genomic DNA was isolated from bulk transfectant cultures and from the resulting clonal lines using the QIAamp Blood Mini Kit (Qiagen, Hilden, Germany). To detect the presence of *hdhfr* integration at the GOI, PCR was performed upon isolated gDNA using the Phusion High-Fidelity PCR Mastermix (New England BioLabs, Ipswich, MA), according to the manufacturer’s instructions. For screening purposes, PCR reactions were also performed direct-from-culture, using a 1:100 dilution of parasitized packed human erythrocytes in the final PCR reaction.

Parasites that were wild type at the *plasmepsin II* locus were detected using primers located upstream and downstream of the gene (PM2 5’ Upstream Fwd and PM2 3’ Downstream Rev), respectively. Integration of the *hdhfr* cassette at the *plasmepsin II* locus was detected using a forward primer upstream of the gene and a reverse primer located in the *hdhfr* cassette (PM2 5’ Upstream Fwd and hdhfr cassette 5’ Rev). To detect parasites that were wild type at the *plasmepsin III* locus, a forward primer specific for the sequence between the homology regions and a reverse primer located downstream of the gene (PM3 Btwn HRs Fwd and PM3 3’ Downstream Rev) were used. Integration of the *hdhfr* cassette at the *plasmepsin III* locus was detected using a forward primer found in the *hdhfr* cassette and a reverse primer located downstream of the gene (*hdhfr* cassette 3’ Fwd and PM3 3’ Downstream Rev). All *plasmepsin II/III* KO transfectants were probed for integration at both the *plasmepsin II* and *plasmepsin III* loci. The presence of plasmid was probed using primers specific to the backbone of the pGEM*hdhfr+* template vector (pGEM_hdhfr+ Fwd and pGEM_hdhfr+ Rev) for all transfections. All primer sequences are listed in S8 Table.

### Quantitative PCR

To determine copy numbers of *pfmdr*1, *plasmepsin* II, and III, qPCR was performed on genomic DNA (extracted with QIAmp Blood Mini Kit, Qiagen, Hilden, Germany) as previously described(24) with the following modifications: amplification reactions were done in MicroAmp 384-well plates in 10 μl SYBR Green master mix (Applied Biosystems, Foster City, CA), 150 nM of each forward and reverse primer and 0.4 ng template. Forty cycles were performed in the Applied Biosystems ViiA^TM^ 7 Real-time PCR system (Life Technologies, Carlsbad, CA). *pfmdr1* primers were designed after Price, et al.(67) whereas *β- tubulin* primers for the endogenous control were designed after Ribacke, et al.(68). To test for integration of the *hdhfr* cassette into the *plasmepsin* locus, two primer sets were designed locating between the homology regions for *plasmepsin II* (RTPCR PMII forw and rev) and *III* (RTPCR PMIII forw and rev) as well as on the *hdhfr* gene (*hdhfr*_RTPCR_F and R). Technical replicates were run in quadruplicates. Copy numbers were considered increased (> 1) when the average of three biological replicates was above 1.6. Primers are listed in S8 Table.

### Southern Blotting

Parasite-infected red blood cells were lysed with 0.15% saponin (Sigma-Aldrich, St. Louis, MO) in PBS and gDNA was isolated from freed parasites via phenol-chloroform extraction. For each clone, 5 ug gDNA was digested with the following restriction enzymes: *Eco*O1091 and *Xmn*I for the PMII KO clones; *Kpn*I and *Nsi*I for the PMIII KO clones; *Eco*O1091, *Xmn*I, *Kpn*I and *Nsi*I for the PMII/III KO clones; *Afl*II and *Nci*l for the PMII KO and PMIII KO clones in a G8 parental background and all PMII/III KO clones. As controls, gDNA from the parental lines (KH001_053_G8_ and KH001_053_G10_) and corresponding plasmid vectors were also digested. Digested gDNA was resolved on a 1% agarose gel in TAE and transferred to Amersham Hybond – N^+^ nylon transfer membrane (GE Healthcare, Chicago, IL). Probe hybridization and horseradish peroxidase-mediated signal detection were performed using the ECL Direct Nucleic Acid Labeling and Detection System (GE Healthcare, Chicago, IL), following the manufacturer’s instructions. DNA probes were designed to detect *hdhfr*, PMII 3’ HR, or PMIII 3’ HR. The *hdhfr* probe was PCR amplified from plasmid vector, while the PMII 3’ HR and PMIII 3’ HR probes were PCR amplified from KH001_053_G10_ gDNA; the corresponding primers are listed in S8 Table.

### *In vitro* drug susceptibility assays by SYBR Green I staining

Drug susceptibility assays were performed using the SYBR Green I method as previously described(69). In brief, tightly synchronized 0-6 h post-invasion rings at 1% parasitemia and 1% hematocrit in 40 uL of 0.5% Albumax-complimented media were grown for 84h in 384-well clear bottom plates, in the presence of different drug concentrations. Drug assays were extended from the standard 72h drug exposure to 84h due to the tight synchronization of the parasites. All drug conditions were performed in three technical replicates, with at least three biological replicates. Drugs were dispensed in 24-point dilution series of PPQ (0.07nM-50uM) and 12-point dilution series of all other drugs (CQ, E64, MQ, DHA, WR99210, Pepstatin A Methyl Ester, and AQ; Sigma-Aldrich, St. Louis, MO) into 384-well plates by an HP D300e Digital Dispenser (Hewlett Packard, Palo Alto, CA). Growth at 84 h was quantified by staining parasite DNA with SYBR Green I (Lonza, Visp, Switzerland) for 24h, and measuring relative fluorescence units at an excitation of 494 nm and an emission of 530 nm using a SpectraMax M5 (Molecular Devices, San Jose, CA). EC_50_ values, for all drugs except PPQ, were calculated using a nonlinear regression with the log(inhibitor) vs. response-Variable slope curve-fitting algorithm using GraphPad Prism version 8-10 (GraphPad Software, La Jolla, CA). PPQ susceptibility was quantified using Area Under the Curve (AUC) between the two local minima of the high-does peak, as previously described (24).

PPQ stocks were resuspended in a 0.5% lactic acid/0.1% Triton X-100 aqueous solution, CQ stocks were resuspended in a 0.1% Triton X-100 aqueous solution, and all other drugs were resuspended in dimethyl sulfoxide (DMSO).

To test the effects of DV stress in modulating PPQ response, parasites were treated with a 24- point dilution series of PPQ in combination with a fixed concentration of the additional inhibitor (E64: 2.6 uM or 3.9 uM, Pep A ME: 5 uM or 7.5 uM, CCCP: 5 uM or 15 uM, and Con A: 0.1 nM or 0.2 nM or DMSO as a control. Relative growth was normalized in comparison to parasite growth in the presence of E64, Pep A ME, CCCP, or Con A at the test concentration alone.

To determine the role of pH in drug susceptibility, parasites were exposed to a dilution series of PPQ or CQ in pH-adjusted Albumax-complimented media for 84h at pH values of ∼6.75, ∼7.5, or ∼8.25

### Cellular heme fractionation assay

Baseline levels of different heme species in the parasite lines were determined using pyridine-based detergent-mediated cellular heme fractionation assays described in detail by Combrinck et al. (38, 39) For the drug exposure experiments with KH001_053_G10_ and KH001_053_G8,_ parasites were synchronized to ring stages using two to three cycles of sorbitol treatment and early rings were incubated at 37°C at 5% parasitemia and 2% hematocrit in 24-well plates. After 32h, late trophozoites were harvested by lysing RBCs with 0.05% saponin followed by multiple washes with 1×PBS (pH 7.5). Pellets were then resuspended in 1×PBS and stored at −80 before further analysis. An aliquot of the trophozoite suspension was stained by SYBR Green I and quantified via flow cytometry (as per above) to determine the total number of trophozoites. For determining the heme composition throughout the life cycle, we increased the starting volume to 10ml of 5% parasitemia at 2.5% hematocrit and further tightened the synchronization by a percoll gradient and sorbitol synchronization 6h later. Parasites were harvested at 24, 36 and 42h post sorbitol synchronization. One set of parasites was exposed to 2μM PPQ between 24 and 36h and harvested at 36h.

Samples were thawed and DV content released from trophozoites by hypotonic lysis and sonication (53 kHz, 320 W). Parasite fractions corresponding to digested Hb, free heme-Fe and Hz were then carefully recovered through centrifugation and treatment with HEPES buffer (pH 7.4), 4% SDS, 25% pyridine solution, 0.3M HCl and 0.3M NaOH.

The UV-visible spectrum of each heme fraction was measured as a Fe3 +-heme-pyridine complex using a multi-well SpectraMax M5 plate reader (Molecular Devices, San Jose, CA). The total amount of each heme-Fe species in a sample was quantified using a heme standard curve and interpolation. The percentage of each species per sample was used to compare between the different strains and conditions. Two-tailed unpaired student’s t tests were used for comparing PPQ treated vs untreated lines at the same time point or between KH001_053_G10_ and the other strains.

### Flow Cytometry to Quantify Parasitemia and stage

Parasites were stained in 10X SYBR Green I in 1X PBS for 30 minutes in the dark at 37°C. The staining solution was removed, and cells were resuspended in five times the volume of the initial volume of PBS. FACS data acquisition was performed on a MACSQuant VYB (Milteni Biotec) with a 488 nm laser and a 525 nm filter and analyzed with FlowJo 2. RBCs were gated on the forward light scatter and side scatter and infected RBCs were detected in channel B1. At least 100,000 events were analyzed per sample. Parasites were considered trophozoites when the DNA content was 3 times higher than the ring stage signal.

### Digestive Vacuole pH determination

Saponin-isolated trophozoite-stage parasites containing the membrane-impermeant pH-sensitive fluorescent indicator fluorescein-dextran (10,000 MW; Invitrogen) in their DVs were prepared as outlined previously (48, 51). The isolated parasites were washed and suspended in malaria saline (125 mM NaCl, 5 mM KCl, 1 mM MgCl_2_, 20 mM glucose, 25 mM HEPES; pH 7.1) at a density of ∼2–3 × 10^7^ cells/ml. 100 uL of cell suspension was added to an equal volume of malaria saline containing test compounds at 2x test concentrations in a 96-well clear-bottomed black plate. The pH of the DV was monitored at 37°C over 1.5h using a Molecular Devices M5i plate reader (excitation 490 and 450 nm, emission 520 nm). Fluorescence ratios (490/450) were calibrated to pH units using calibration buffers at pH 4.5, 5.1, 5.7 and 6.3 (130 mM KCl, 1 mM MgCl_2_, 20 mM glucose, 25 mM HEPES; 180 nM Nigericin, 100 nM Concanamycin A) as described previously (48, 51).

## Acknowledgments

DFW received funding from the ExxonMobil Foundation.

## Supporting information captions

**S1 Figure. Confirmation of *plasmepsin II* and *III* single KOs clones by Southern blot.**

A) Schema of original loci, homology plasmids used for integration into the locus and resulting edited loci. Restriction enzyme sites and expected band sizes for Southern blots are indicated in the schema in orange for *plasmepsin II* and in blue for *plasmepsin III* KOs. B) Southern blots with different probes, expected band sizes are indicated by arrows. Clones indicated with a star were used for phenotyping. Plasmids were either transfected in circular (cir) or linearized (lin) form. C) WR99210 targets primarily the plasmodial *dhfr* and an increase in EC_50_ correlated with the presence of one or several hdhfr cassettes present. Shown is the average EC_50_ and standard deviations of 3 biological replicates for each clone (one-way ANOVA with Dunnet post-test compared to 1D with a single integration of the *hdhfr* cassette. **** P < 0.0001).

**S2 Figure. Confirmation of *plasmepsin II*/*III* double KOs clones by Southern blot.**

A) Schema of original parental loci, homology plasmid used for integration into the loci and resulting edited locus which is identical for both parents. Restriction enzyme sites and expected band sizes for Southern blots are indicated in the schema in orange for plasmepsin II and in blue for plasmepsin III KOs. B) Southern blots with plasmepsin III probe, expected band sizes are indicated by arrows. Clones indicated with a star were used for phenotyping. C) Average EC_50_ and standard deviations of 3 biological replicates of WR99210 for each clone except 4E (1N). Nd: not determined

**S3 Figure. Confirmation of *plasmepsin II*/*III* double KOs clones by Southern blot.**

A) Schema of original parental loci, homology plasmid used for integration into the loci and resulting edited locus which is identical for both parents. Restriction enzyme sites outside the locus were selected to confirm complete deletion of the regions between the homology regions and expected band sizes for Southern blots are indicated in the schema. Clones indicated with a star were used for phenotyping. B) The same Southern blots was hybridized three times with the plasmepsin II, plasmepsin III or the hdhfr probe. Expected band size for each probe are indicated with arrows. The loss of hybridization for the plasmepsin II probe confirms the deletion and fusion of plasmepsin II and plasmepsin III in the KO clones.

**S4 Figure. Parasite growth in different concentrations of drug compared to no drug control.**

Ring stage parasites were either grown in complete media or in complete media with the addition of CCCP (15 or 5 μM), E64 (3.9 or 2.6μM) Pepstatin A (7.5 of 5μM), Concanamycin (0.1 or 0.2nM), 10μM DHA (dead) or 0.5% DMSO for 72h. Growth was measured by the incorporation of SYBRGreen into DNA, read by a spectrometer and normalized to parasites grown in media only.

**S5 Figure. Effect of protease inhibitors on PPQ efficacy in parasites with a single (KH001_053_G10_), double (KH001_053_G8_) or multiple (KH004_057) plasmepsin loci.**

Parasites were exposed to increasing levels of PPQ in the presence of DMSO or A) E64 at either 2.6μM or 3.9μM concentrations or B) pepstatin A methylether (Pep A ME) at either 5μM or 7.5μM concentration. Shown is one example of 3 biologically independent experiments run in triplicates. C) Average and SD of the area under the curve (AUC) between the local minima for three biological replicates. No statistically significant difference was detected between PPQ alone and combination of E64 or PepA by ordinary one-way ANOVA with Tukey posttest.

**S6 Figure. Effect of internal or external pH on CQ efficacy.**

A-F) Parasites were exposed to increasing levels of CQ in the presence of DMSO or CCCP at either 5 μM or 15μM concentrations. G-L) Parasites were exposed to increasing levels of CQ in acidic (pH6.74), normal (pH7.5) or basic (pH8.24) media. Shown is one example for each line tested of three biologically independent experiments run in triplicates. The EC_50_ was calculated where possible (an # indicates when parasites were not killed completely at the highest concentration), and the average and SD are shown in (F and L). Statistics show one-way ANOVA with Tukey post-test for each strain tested in the presence of CCCP or two tailed paired Student’s t test for the external pH changes: *, P < 0.05; **, P < 0.01, and ***, P < 0.001.

**S1 Table. Analysis of parasite clones from different transfections to confirm proper integration into the genome by PCR and qPCR.** Plasmids were either transfected in circular form or were linearized with BglI before transfection (lin). Shown are all the clones analyzed for every transfection that was recovered. Integration was confirmed by PCR of the 5’ and 3’ region and appearance of a band at the right size was considered a positive result (green check mark). The absence of the transfection plasmid was screened for by using primers targeting the backbone of the plasmid which is lost after correct integration (red cross). Quantitative PCR was also used to confirm copy numbers of *plasmepsin II* and *III* as well as the *hdhfr* gene inserted into the locus. Primers are listed in S8 Table.

**S2 Table. EC_50_ data for various drugs tested.** Parasite lines were exposed to various concentration of drug and the EC_50_ was calculated for each drug. Each experiment was run in triplicate and at least 3 biological replicates were performed for each parasite line/clone. Shown are the EC_50_, SD and number for each line.

**S3 Table. Combination assays results (Fig 5E-L and S6 Fig).** A) Combination assays of parasites in the presence of increasing concentrations of PPQ and a constant concentration of E64, PepA, Con A or CCCP measured as AUC. B) EC_50_ for ConA and CCCP. C) Combination assays in the presence of increasing concentrations of CQ and a constant concentration of CCCP measured as EC_50_.

**S4 Table. Data for heme fractionation of PPQ treated and untreated parasites with variable plasmepsin copy numbers (Fig 3A-C).** Different heme species were extracted from parasites by subsequent cellular fractionation steps. Tightly synchronized KH001_053_G10_ and KH001_053_G10_ ring stage parasites were exposed to various PPQ concentrations for 32h. The amount of iron species (Fe) per sample was estimated based on the standard curve run for each biological replicate and the percentages for each species per sample was calculated. Shown are each value from three biological replicates run in quadruplicates with the average and SD of percentage of hemozoin Fe, hemoglobin, and free heme Fe. Statistical comparisons of the drug-treated lines to their untreated controls were performed using two-tailed unpaired student’s t tests *, p < 0.05; **, p < 0.01.

**S5 Table. Data for heme fractionation of Plasmepsin KO parasites throughout the life cycle and exposed to PPQ. (Fig 3D-F and Fig 4).** Tightly synchronized parasites were harvested at different time-points throughout the life cycle and average and SD of percentage of hemozoin Fe, hemoglobin, and free heme Fe are shown for three independent experiments for parasites with a single locus (KH001_053_G10_), duplicated locus (and KH001_053_G8_), G8_pmII_KO_, G8_pmIII_KO_, or G8_pmII/III_KO_. Statistical comparisons at each time point were performed between the single copy KH001_053_G10_ parasite and all other lines using two-tailed unpaired student’s t tests *, p < 0.05; **, p < 0.01. Additionally, parasites were incubated with 2μM PPQ from 24 to 36h post synchronization and harvested at 36h. Statistical comparisons of treated to untreated parasites at 36h timepoint from three independent biological replicates were performed using two-tailed unpaired student’s t tests *, p < 0.05; **, p < 0.01, p < 0.01, *** p > 0.001.

**S6 Table. Effect of extracellular pH on drug response (Fig 5A and S6 Fig).** Parasites were exposed to increasing levels of CQ or DHA for 72h or PPQ for 84h in acidic (pH6.74), normal (pH7.5) or basic (pH8.24) media. Included are the EC_50_ or AUC data for four biologically independent experiments run in triplicates for Dd2, 3D7, KH001_053_G10_, KH001_053_G8_ and KH004_057 including the average and SD of the AUC or EC_50_. Unpaired student’s t test between pH at 7.5 and lower or higher pH if more than 3 values could be determined **, P < 0.01, ***, p<0.001,

**S7 Table. Digestive vacuole pH measurement of fluorescein-dextran loaded Dd2 parasites.**

A) DV pH traces of Dd2 parasites exposed to PPQ (50 μM, 10 μM, 5 μM and 1 μM) and control treatments Concanamycin A (ConA, 100 nM), CCCP (10 µM), CQ (10 µM) and NH_4_Cl (10 mM) over 90 minutes run in technical duplicates. B) DV pH was quantified for each treatment as an average of the measurements taken between 45 and 60 mins of compound. Shown is the average pH for 3 to 4 independent experiments (performed with blood from different donors).

**S8 Table. List of primers used.** This table includes all primers used in this study.

